# *In vivo* homopropargylglycine incorporation enables nascent protein tagging, isolation and characterisation from *Arabidopsis thaliana*

**DOI:** 10.1101/2021.03.04.433937

**Authors:** Nathan D. Tivendale, Ricarda Fenske, Owen Duncan, A. Harvey Millar

## Abstract

Determining which proteins are actively synthesised at a given point in time and extracting them for analysis is important to understand plant responses. Here we show that the methionine (Met) analogue homopropargylglycine (HPG) enables BONCAT (Bio-Orthogonal Non-Canonical Amino acid Tagging) of proteins being synthesised in Arabidopsis plants or cell cultures, facilitating their click-chemistry enrichment for analysis. The sites of HPG incorporation could be confirmed by peptide mass spectrometry at Met-sites throughout protein AA sequences and correlation with independent studies of protein labelling with ^15^N verified the data. We provide evidence that HPG-based BONCAT tags nascent plant proteins more efficiently than azidohomoalanine (AHA)-based BONCAT in Arabidopsis and show that AHA’s induction of Met metabolism and greater inhibition of cell growth rate than HPG likely limits AHA incorporation at Met sites in Arabidopsis. We show HPG-based BONCAT provides a verifiable method for determining which plant proteins are being synthesised at a given time point and enriches new protein molecules from the bulk protein pool for identification, quantitation and subsequent biochemical analysis. Enriched nascent polypeptides were found to contain significantly fewer common post-translationally modified residues than the same proteins from whole plant extracts, providing evidence for age-related accumulation of PTMs in plants.

## Introduction

In all cellular organisms, protein pools are continually being renewed through a cyclic process of synthesis, degradation and re-synthesis of protein molecules. Identifying and quantifying which proteins are being synthesised at a given point in time is complicated, especially if synthesis replaces proteins that are degraded, leading to no net change in protein abundance. Traditionally, [^35^S]Met labelling has been used to label proteins before visualisation of protein synthesis was achieved through gel electrophoresis and autoradiography (Geiken et al., 1998). However, this approach has limitations in its ability to identify labelled proteins and renders them radioactive. Non-radiolabelled methods based on stable isotope labelling have enabled large scale measurement of protein synthesis: for example SILAC (Stable Isotope Labelling with Amino acids in Cell culture), fertilizing with ^15^N based salts or growing in air supplemented with ^13^CO_2_ (Lewandowska et al., 2013; Bagert et al., 2014; Nelson et al., 2014; Ishihara et al., 2015; Nelson and Millar, 2015; Fan et al., 2016; Li et al., 2017). These methods rely on peptide mass spectrometry to identify old and new protein populations but do not enable nascent proteins to be isolated, which would increase the potential sensitivity of identification and quantification and allow for subsequent biochemical analysis of enriched samples of nascent proteins.

Bioorthogonal noncanonical amino acid tagging (BONCAT) with click-chemistry enabled amino acid analogues (Dieterich et al., 2006) that harbour azide and alkyne groups can facilitate the enrichment of tagged proteins. An effective BONCAT strategy relies on supplying noncanonical amino acids (ncAAs) that have sufficient homology with natural amino acids to be incorporated into an organism’s proteins. The selectivity of aminoacyl-tRNA synthetases in charging tRNA with ncAAs, binding of ncAA-tRNAs at the A site between ribosome subunits, and steric hindrance in the emerging and folding of the ncAA-containing nascent polypeptide chains will dictate the suitability of different click chemistry-enabled ncAAs in different organisms. Azidohomoalanine (AHA) and homopropargylglycine (HPG) are Met analogues that have been shown to be efficiently incorporated into newly-synthesised proteins in a variety of systems, including mammalian cell culture (Dieterich et al., 2006; Bagert et al., 2014), mice (McClatchy et al., 2015), *Drosophila melanogaster* (Erdmann et al., 2015), *E. coli* (Kramer et al., 2011) and cell-free systems (Worst et al., 2015; Gao et al., 2019). While methionyl-tRNA synthetase inefficiently charges the Met-tRNA with AHA and HPG, when present in excess to Met they can make sufficient L-AHA/HPG-tRNA to effectively label nascent proteins (McClatchy et al., 2015; Calve et al., 2016). Met has been chosen as it is a relatively rare amino acid (Gilis et al., 2001) in proteins, ensuring that modified proteins still fold and locate appropriately within cells (Calve et al., 2016; Zanobini et al., 2018). As AHA and HPG tagged proteins can be enriched by azide-alkyne cycloaddition (‘click chemistry’), lower levels of tagging are detectable and hence shorter experimental timeframes for protein synthesis studies are achievable.

For many years, while trace amounts of [^35^S]Met were supplied to plants for protein synthesis studies (e.g. Cordewener et al., 1994; Geiken et al., 1998), BONCAT was not applied. This was at least in part because Met (and by extension Met analogues) supplied in excess to plant cell culture have been shown to inhibit growth, even at concentrations as low as 100 μM (Vakkari, 1980) and accumulation of Met in whole plant tissues results in stunted plant growth (Boerjan et al., 1994). Unlike in mammalian cells, where Met is an essential nutrient and promotes cell growth, plant cells make their own Met and this is involved in numerous biochemical processes. In fact, studies of the metabolic fate of Met in *Lemna paucicostata* indicate that only about 20 % of Met is used in protein synthesis in plants (reviewed in Giovanelli, 1987). For proper cellular functioning, it is important for plants to maintain consistent levels of S-adenosyl-methionine (SAM). Plants keep the level of SAM constant through SAM-dependent methylation of Met to produce *S*-methylmethionine (SMM; Rébeillé et al., 2006). Thus, plant cells accumulate SMM, while maintaining steady concentrations of SAM, to cope with excess Met (Rébeillé et al., 2006).

The absence of BONCAT studies in plants changed when it was reported that AHA could be incorporated into nascent proteins in whole Arabidopsis plants *via* seedling flood (Glenn et al., 2017). One report of protein tagging with AHA in plants preceded this report, but no data were shown and no details of the method were given (Echevarría-Zomeño et al., 2016). Glenn et al. (2017) provided a detailed methodological report showing AHA incorporation into proteins was higher under plant heat stress and supported this claim using AHA-specific antibodies; they also reported that nascent Arabidopsis proteins could be enriched by click chemistry-enabled affinity chromatography and enriched proteins identified by LC-MS/MS. This has promoted the potential of BONCAT in plants and led to a series of reports noting the potential for AHA labelling to be extended to these organisms (e.g. Sesma et al., 2017; Lee and Bailey-Serres, 2018; Mazzoni-Putman and Stepanova, 2018; Hatzenpichler et al., 2020; Tivendale et al., 2020; Yu et al., 2020).

Here, we have followed up on these reports to better understand how AHA impacts plant growth, to clarify which proteins are tagged by AHA and to compare AHA with the alternative BONCAT reagent HPG and also to ^15^N-labelling of endogenous amino acid pools to more fully understand the utility of BONCAT for plant science. We conclude that BONCAT is a useful tool for study of proteins in plants but HPG has better utility for labelling and isolation of nascent plant proteins for a variety of applications in the general analysis of plant protein synthesis, as well as for identification and/or isolation of tagged proteins for independent study.

## Results

### HPG is more effective than AHA for protein tagging in whole plants

Whole Arabidopsis plants treated with AHA *via* seedling flood, have been previously reported to show incorporation of AHA into proteins using specific antibodies, and separately, proteins enriched using click chemistry have been identified using peptide mass spectrometry (MS) (Glenn et al., 2017; Yu et al., 2020). While these studies show the general utility of BONCAT, neither provided MS/MS evidence of AHA residues replacing Met residues in tagged proteins identified by MS. To determine the extent of AHA labelling in AHA-treated plants, we treated Arabidopsis plants with AHA using the same technique. Proteins isolated from plants treated with AHA had MS-confirmed AHA tags in only one out of the 3022 protein groups (0.03 %; 1 tagged peptide was identified) in total extracts of whole seedlings (Figure 1A, Supplemental Table S2). As an independent comparison we also treated plants with the alternative ncAA, HPG, using the same method and, in contrast, these plants displayed HPG tags in 11 out of 2917 protein groups (0.37 %; 16 unique tagged peptides were identified; Figure 1A, Supplemental Table S3). Therefore, HPG appeared to be more effective than AHA for tagging nascent proteins in whole plants, although the seedling flood in our hands appeared to have limited use for studying protein synthesis in whole plants. Assessment of green leaf area viewed from above showed that flooding with AHA and HPG negatively affected plant increase in green area compared to the mock, however the worst affected were those treated with AHA (Figure 1C). To further investigate the potential use of AHA and HPG as nascent-protein-tagging agents in plants, we therefore turned to a series of experiments using Arabidopsis cell culture where a flooded environment is the control state of cells.

**Figure 1.**
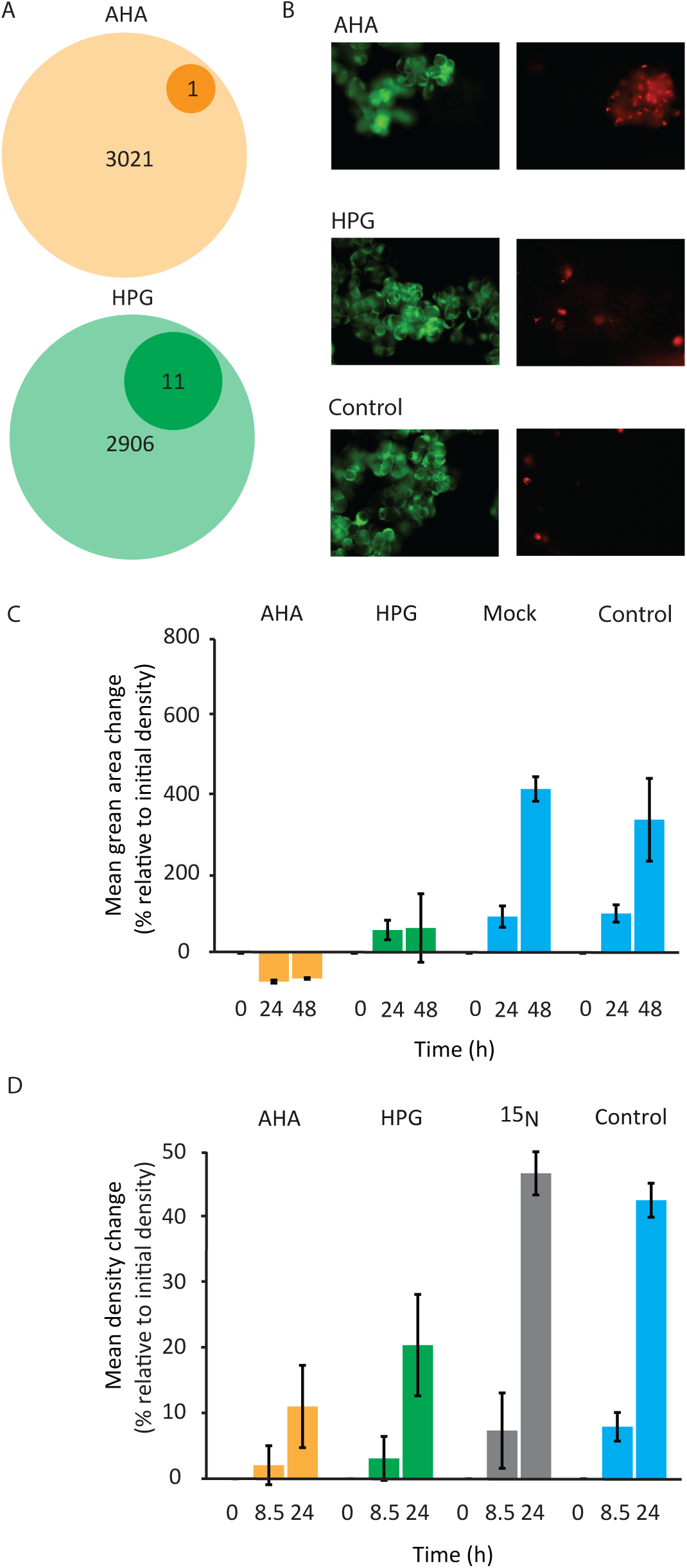
The impact of non-canonical amino acid labelling on Arabidopsis cell culture and whole plants. (A) AHA and HPG tagging of Arabidopsis leaf proteome. (B) Viability staining of Arabidopsis cell cultures. A combination of propidium iodide and fluorescein diacetate was used. Red fluorescing cells are dead and green fluorescing cells are alive. (C) Effects of treatment with AHA or HPG, *via* seedling flood, on whole Arabidopsis plant growth, in terms of green area in images. Mean (n = 2) ± se for each time point are shown. (D) Cell density changes of Arabidopsis cell cultures treated with various chemicals. Mean (n = 3) ± se for each time point are shown.

### The effect of Met, AHA, HPG and ^15^N labelling on plant cell growth

To begin a systematic comparison of BONCAT labelling in plant cell culture, we examined the effects of AHA, HPG and ^15^N on cell culture growth. Met and the ncAAs were effectively taken up into Arabidopsis cells, as determined by targeted LC-MS (Table 1). The recorded cell content of ncAAs correlated with a loss of cell growth rate in AHA-treated cells, but no effect on the growth rate of ^15^N-treated cells was observed; a less severe decrease in growth rate was also observed in HPG-treated cells (Figure 1D). Assessment of cell death showed HPG caused less cell death than AHA in Arabidopsis cell cultures (Figure 1B).

**Table 1.** Entry of Met, AHA and HPG into Arabidopsis cells. Amount of Met, AHA and HPG in Arabidopsis cells after their incubation with 1 mM of each of these compounds for 8 or 24 hours. Absolute abundances were determined by LC-MS. Means ± se (n = 3) are shown.

| Time (h) | Met (mg/g (FW)) | AHA (mg/g (FW)) | HPG (mg/g (FW)) |
| --- | --- | --- | --- |
| 8 | 0.3 ± 0.1 | 0.083 ± 0.003 | 0.075 ± 0.003 |
| 24 | 0.7 ± 0.4 | 0.077 ± 0.002 | 0.084 ± 0.002 |

### AHA and HPG stimulated in-vivo synthesis of Met and related metabolites

To reveal the possible cause of the cell culture growth inhibition caused by AHA, HPG, we determined changes in the levels of Met and four related metabolites in Arabidopsis cell cultures treated with AHA, HPG or Met over the course of 24 h (Figure 2). After 24 h there was no significant difference between the level of *S*-adenosylhomocysteine (AdoHcy), homocysteine (Hcy) or SAM (p > 0.05), but the levels of 5-methylthioadenosine (5-Me-S-Ado) were significantly higher (p < 0.05) in the cells treated with Met and AHA and, to a lesser extent, in cells treated with HPG. The differences in the average level of Met after 24 h appeared different, but the large standard deviation rendered this difference not statistically significant. Nevertheless, the observed disruption of Met metabolism may be the cause of effects on cell growth that we observed in cell cultures treated with AHA and the less dramatic effects observed in cell cultures treated with HPG. After 24 h of treatment, the ratio of the supplied ncAA to endogenous Met was 1.6 for AHA and 4.8 for HPG (Table 1).

**Figure 2.**
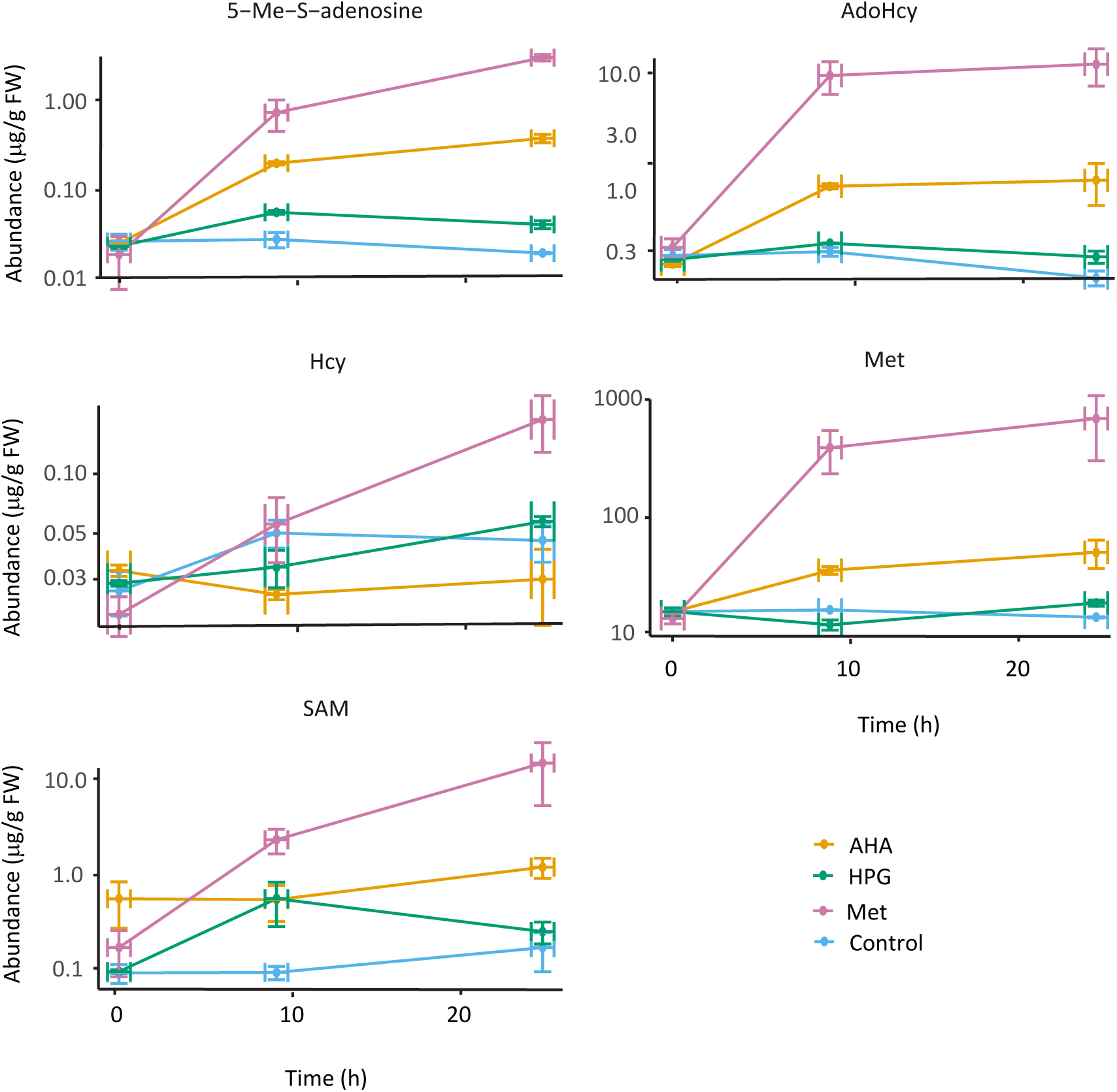
Methionine metabolism in response to non-canonical amino acids labelling in Arabidopsis. Levels of methionine and four methionine-related metabolites in Arabidopsis cell cultures treated with azidohomoalanine (AHA), homopropargylglycine (HPG) or methionine, over 24 hours, as determined by LC-MS.

### HPG was more efficiently incorporated into proteins replacing Met than AHA

After establishing that HPG caused less metabolic disruption that AHA in Arabidopsis cell cultures, we used LC-MS/MS to determine the number of proteins tagged with AHA or HPG after 24 h of exposure. We identified a similar total number of protein groups in AHA- and HPG-treated cells (3177 and 3146 protein groups, respectively) using data-dependent acquisition (DDA) proteome profiling (Supplemental Tables S4 and S5). However, in the case of treatment with AHA, none of the proteins showed any peptide mass spectrometry evidence of AHA replacing Met in the amino acid sequence of identified peptides. In contrast, 100 of the protein groups identified in the cells treated with HPG (3.2%) showed mass spectrometry evidence of HPG replacing Met in peptides matching their amino acid sequences (Table 2, Figure 3). Thus, it was clear that HPG was a much more effective ncAA for nascent enzyme tagging, not only because it caused less metabolic disruption and less cell death, but also because tagging of proteins could be verified even in whole proteome extracts analysed by DDA. Proteomics datasets from AHA- and HPG-treated cells were inseparable by PCA, but both of these datasets were slightly separated from proteomics datasets from untreated cells, indicating that AHA and HPG treatments alone have some effect on the plant cell proteome composition (Figure 4A).

**Figure 3.**
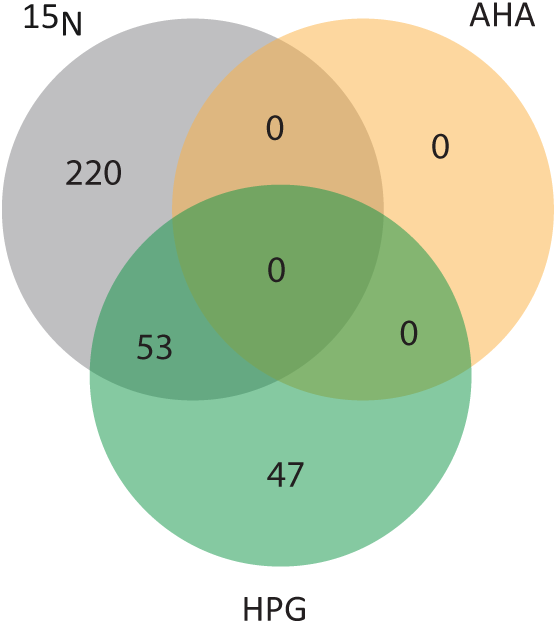
Nascent protein labelling in Arabidopsis cell culture. Number of protein groups in ^15^N-, AHA- and HPG-treated Arabidopsis cell cultures with evidence of ^15^N labelling or replacement of methionine with AHA or HPG.

**Figure 4.**
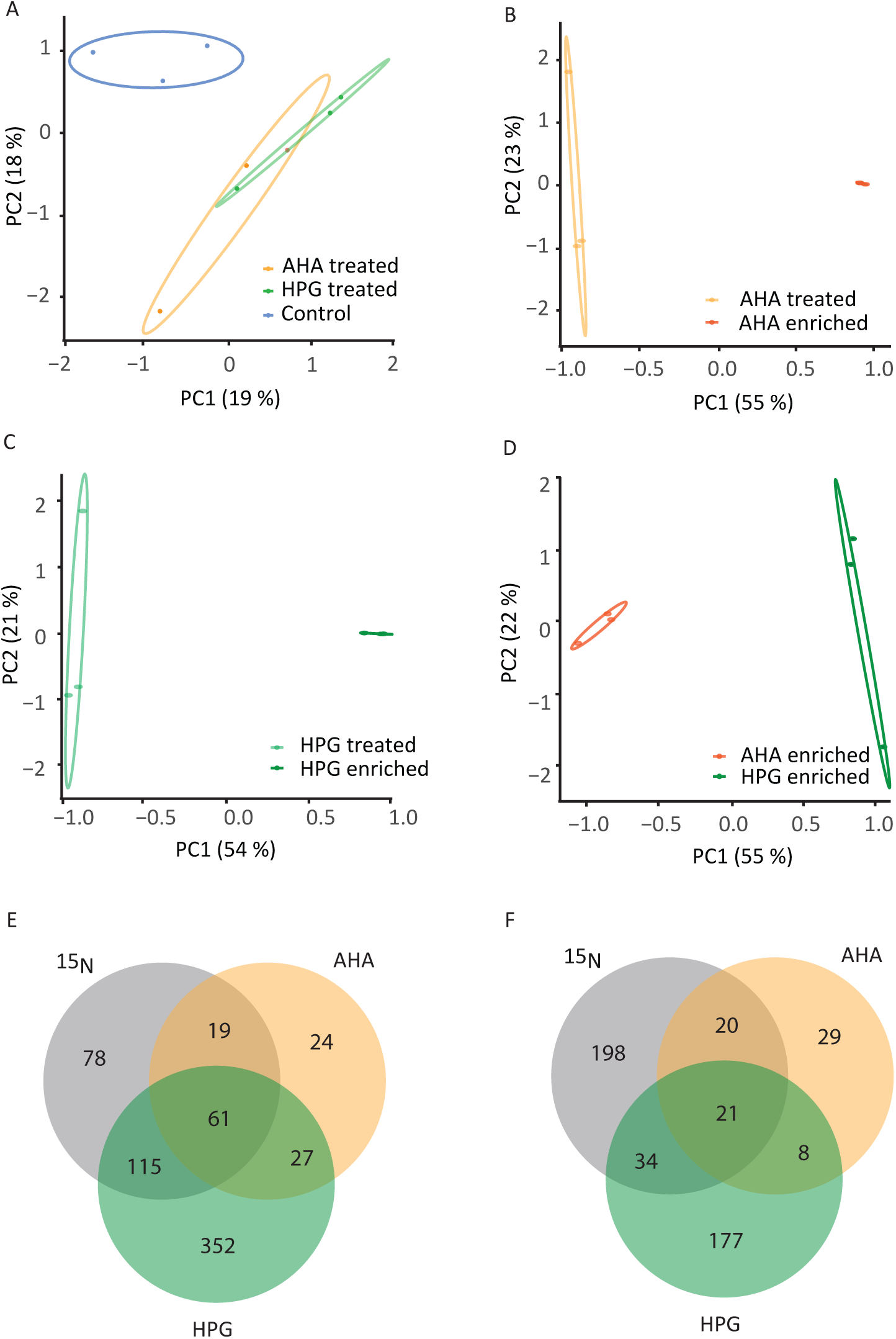
Principal component analysis for Arabidopsis protein sets from labelling and click-chemistry enrichment. Total protein extracts (A), resin-enriched and total protein extracts from cells treated with AHA (B) or HPG (C) and resin-enriched protein extracts from cells treated with AHA or HPG (D). Comparison of the number of protein groups labelled with ^15^N and resin-enriched from Arabidopsis cells treated with AHA or HPG (E). Comparison of the number of protein groups labelled with ^15^N and resin-enriched (relative enrichment value > 0) from Arabidopsis cells treated with AHA or HPG (F).

**Table 2.** Numbers of Arabidopsis protein groups identified in non-canonical amino acid enriched samples. No. tagged protein groups (number of protein groups containing at least one Met-to-AHA or Met-to-HPG peptide in bulk protein extracts), No. protein groups enriched (number of protein groups identified in click chemistry resin chromatography enriched samples), enriched and tagged (number of protein groups both enriched and AHA or HPG tagged), enrichment > 0 (number of proteins with higher relative abundance in enriched than in bulk protein extracts), tagged in enriched set (number of proteins with at least one peptide containing at least one Met-to-AHA or Met-to-HPG observed in enriched protein sets). Proteins were counted when they were observed to fulfil the stated criteria in at least two out of three biological replicates.

| Tag | No. tagged protein groups | Sample type | No. protein groups enriched | enriched and tagged | enrichment > 0 | tagged and enrichment > 0 | tagged in enriched set |
| --- | --- | --- | --- | --- | --- | --- | --- |
| AHA | 0 | Digested on resin | 131 | 0 | 78 | 0 | 0 |
|  |  | Cleaved from resin | 47 | 0 | 35 | 0 | 0 |
|  |  | Peptides cleaved from resin after on-resin digestion | 2 | 0 | 2 | 0 | 0 |
| HPG | 100 | Digested on resin | 555 | 74 | 240 | 39 | 7 |
|  |  | Cleaved from resin | 17 | 3 | 14 | 3 | 1 |
|  |  | Peptides cleaved from resin after on-resin digestion | 24 | 8 | 17 | 8 | 2 |

### Enrichment of AHA and HPG-tagged proteins from plant cells

When AHA-tagged proteins from plants were previously reported (Glenn et al., 2017; Yu et al., 2020), proteins were cleaved off the enrichment resin using trypsin, which could result in the ncAA tags remaining on the resin and thus not appearing in peptide mass spectrometry data. However, this lack of direct evidence of AHA-tagging of proteins also leaves open the possibility that proteins from AHA treated-plants may become enriched through alternative plant-specific and/or yet-to-be-defined chemical interactions with the enrichment media, rather than through azide-alkyne cycloaddition.

To verify the specificity of the enrichment of click-chemistry products in our experiments, we enriched intact proteins from cell cultures treated with HPG and AHA (Table 2, Supplemental Table S4–5). AHA enrichment uses alkyne functionalised resin while HPG enrichment uses azide functionalised resin. PCA showed that the HPG enrichment processes produced protein sets of composition and iBAQ abundances that were distinct from the total protein samples extracted from cell cultures (Figure 4B). In contrast, the enriched and bulk datasets from AHA-treated cell cultures were inseparable by PCA (Figure 4C). Interestingly, PCA also showed that the HPG and AHA enriched datasets were distinct from each other (Figure 4D). Previously, it has been shown that enrichment of untreated proteins by this chromatography approach is negligible as it is performed under very stringent conditions with high urea concentrations used for washing and large dilution volumes that will prevent any significant non-covalent associations from being retained (Zhang et al., 2014).

To determine if the enrichment was selective for proteins with the HPG and AHA tag, we used MS to look for the presence of Met-to-HPG and Met-to-AHA substitutions in peptides from bulk proteins from the ncAA-treated cells and compared them with peptides from proteins that were detected in the protein samples after enrichment. The results of these experiments are summarised in Table 1.

For the cell cultures treated with HPG, 555 protein groups were found in the resin-enriched protein samples, and 461 of these were also found in whole tissue extracts; thus the enriched protein groups represent ca. 15 % of those from whole tissue extracts. Of the 555 enriched protein groups, 74 showed evidence of HPG tagging in the bulk data set (Table 1, Supplemental Table S6). We also calculated an relative enrichment value ([riBAQ enriched – riBAQ total]/[riBAQ enriched + riBAQ total]); a value > 0 indicates a protein with a higher relative abundance in the enriched than the total extract. Within the 555 enriched protein groups, 240 (43 %) had an enrichment value greater than 0 and of these, 39 showed evidence of HPG tagging in the bulk data set (Table 1, Supplemental Table S6). It should be noted that not all proteins will produce a peptide with an ncAA that is also detectable by LC-MS/MS, so one would expect that proteins for which no tag can be experimentally detected could increase in abundance during the enrichment process. Proteins with enrichment values less than 0 may be due to a low stoichiometry of HPG tags or weak binding of a non-tagged protein. Only seven of the 555 protein groups showed experimental evidence of HPG tagging in the resin-enriched protein samples, but this could be explained by cleavage of proteins form the enrichment resin with trypsin, leaving many HPG tagged peptides still attached to the resin.

In comparison, for cell cultures treated with AHA, only 131 protein groups were found in the resin-enriched protein samples. Of these, 78 protein groups (60 %) had an enrichment value greater than 0, even though there was no evidence of AHA tagging of peptides for any of these proteins in the bulk or resin-enriched datasets. Comparing the absolute numbers of protein groups in the enriched datasets from both treatments might suggest that AHA is more selectively incorporated into nascent proteins than HPG. However, comparing the percentage of the enriched proteins with enrichment values greater than 0 between the two treatments suggests that AHA may be incorporated to a greater extent than HPG in the proteins that it tags. Notwithstanding the inefficiency of the method, the trends described above were also seen in experiments where the proteins were cleaved from the resin using hydrazine and where peptides that remained bound after on-resin digestion were cleaved from the resin using hydrazine and then separately analysed (Table 2).

Taken together, these results suggest that the click-chemistry enrichment process is somewhat opaque and proteins that are bound through interactions other than azide-alkyne cycloaddition with ncAAs may be contributing to the protein sets identified. As a result we caution against the notion that the presence of a given protein in a click-chemistry enriched protein sample is sufficient to conclude that this protein has been tagged with ncAAs. Rather, enrichment should be confirmed by MS evidence of ncAA incorporation in peptides before or after enrichment to verify members of a nascent protein set. Further, the data from our HPG-treatment experiment shows that not all tagged proteins bind effectively to the resin (our data show that 26 of the 100 Met-to-HPG tagged proteins from the HPG-treated cells were not identified in the alkyne-resin enriched dataset), so there appears to be some unidentified bias in the enrichment process that causes the loss of some bona fide tagged proteins.

### AHA and HPG tag some of the same proteins as ^15^N

To verify how well the ncAA tagged proteins represented the nascent protein pool, we compared them to proteins labelled by incorporation of ^15^N salts into proteins over 24 hours. Out of the 100 protein groups that were found tagged with HPG, 53 were also found in the 273 protein groups that were significantly labelled with ^15^N in a 24 hour period in cell cultures (Figure 3). To order the ^15^N labelled proteins according to their abundance of nascent protein, we multiplied the ^15^N labelled protein fraction (LPF) by the abundance of each protein from untreated cell culture datasets. This abundance was estimated using intensity-based absolute quantification (iBAQ). When we sorted the protein groups list from most to least newly-synthesised protein we found that 35 out of the top 40 protein groups in this list were also verified as HPG tagged (16) and/or resin-enriched from HPG treated samples (32) (Figure 5, Supplemental Table S6). While 27 out of the top 40 protein groups were also resin-enriched from AHA treated samples, none were verified as AHA tagged (Figure 5, Supplemental Table S6).

**Figure 5.**
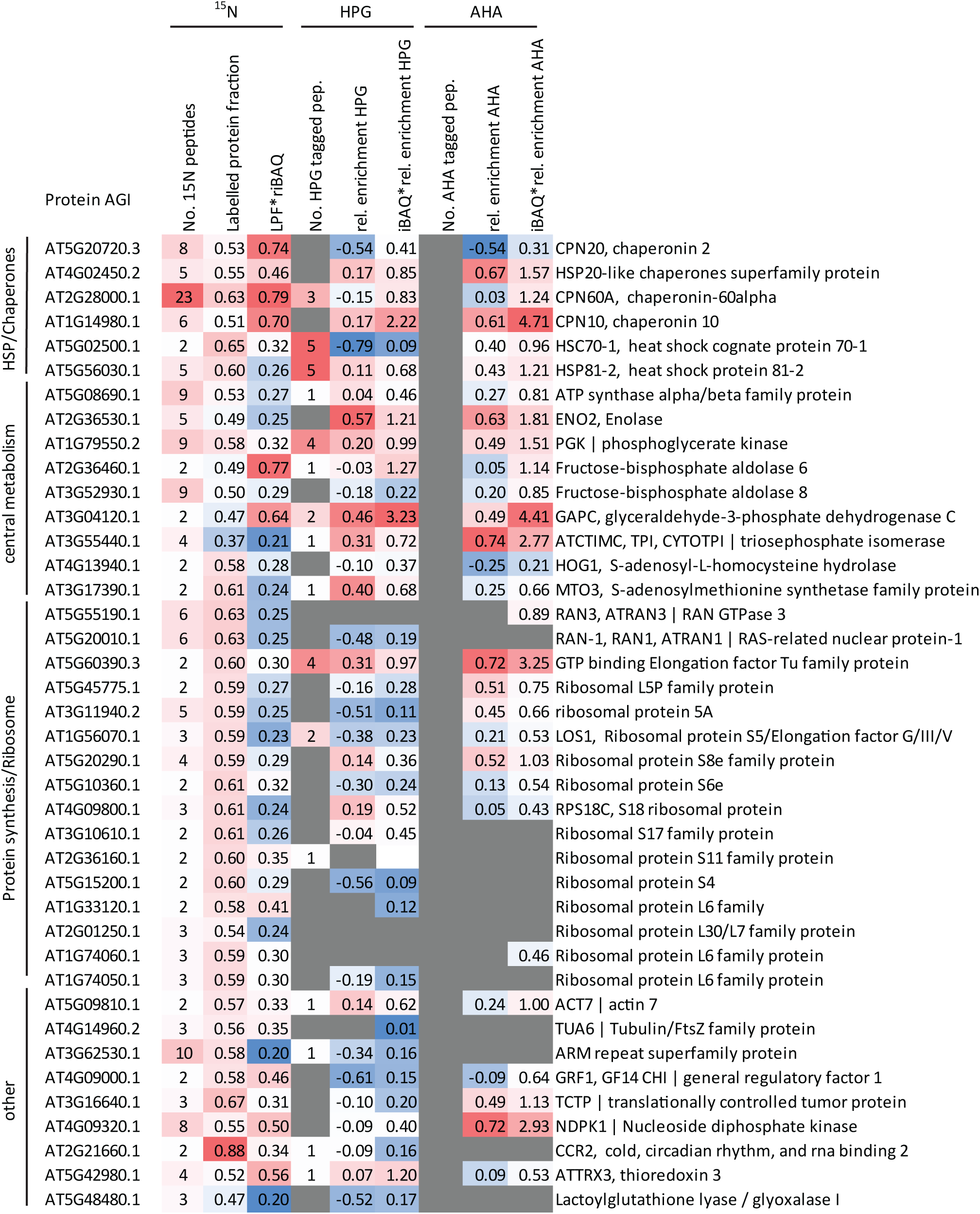
Top 40 newly synthesised Arabidopsis cell culture proteins determined by ^15^N- and non-canonical amino acid labelling. Conditional formatting was applied to each column to highlight differences in values. LPF: labelled protein fraction. riBAQ: relative iBAQ. Protein descriptions are from TAIR10 annotation.

Comparison with labelling by ^15^N provided several avenues to investigate the selectivity of the ncAA enrichment process for nascent proteins. There was a substantial overlap between the protein groups that were labelled with ^15^N and those that were detected in the resin-enriched samples from cells treated with AHA or HPG. Of the 273 protein groups that were tagged with ^15^N, 80 and 176 of these were also enriched in the samples from the cells treated with AHA or HPG, respectively (Figure 4E; 29 % and 64 % of the protein groups labelled with ^15^N were also present after enrichment in samples from cell cultures treated with AHA or HPG, respectively). However, only 61 protein groups were shared between all three lists (Figure 4E); 19 enriched groups were shared only between the ^15^N and AHA enriched dataset and 115 enriched groups were shared only between the ^15^N and HPG enriched dataset. These overlaps decreased substantially when only protein groups with enrichment values greater than 0 were considered (Figure 4F).

### Enrichment is weakly linked to percent methionine, but not to protein length, the number of tags per protein, relative position of tags or abundance of newly synthesised protein

As AHA or HPG tags proteins in the position of Met residues, it is important to clarify if enrichment was significantly biased by the protein percent Met, length, number of tags or the relative position of tags within amino acid sequences. We looked for correlations between each of these attributes and the enrichment values (see Supplemental Figures S1–S6). The only correlation that was apparent was a weak one between the percentage of Met in a protein and its relative-enrichment (Supplemental Figures S1–S2); this correlation was stronger in the case of cells treated with HPG, compared to those treated with AHA. It might be expected that the proteins displaying the highest enrichment would be those with the highest abundance of newly-synthesised material based on ^15^N labelling; however, we found no such correlation (Supplemental Figures S7–S8) for enrichment using either ncAA tag.

### GO term and PO and GO domain enrichment analysis reveals the types of proteins that are tagged and enriched by ^15^N and ncAAs

Using AgriGO (Du et al., 2010; Tian et al., 2017), we observed some degree of GO term enrichment in tagging and ncAA enrichment sets. HPG and ^15^N both favoured tagging of stimulus response proteins, but for HPG they were mainly heat response proteins while for ^15^N they were mainly cadmium response proteins (Supplemental Figures S9–S10). In terms of click-chemistry enrichment, proteins resin-enriched from AHA-treated cells were those involved in carbohydrate metabolism and stress/stimulus response proteins (Supplemental Figure S11), while stress and heat response proteins were overrepresented in samples resin-enriched from HPG-treated cells (Supplemental Figure S12).

Taken together, these data indicate that stress/stimulus response proteins undergo rapid synthesis in Arabidopsis cell cultures. This is likely not a result of the treatment with ncAAs, since stress/stimulus response proteins were also overrepresented in the ^15^N labelled data set and there is no precedent for thinking ^15^N would cause synthesis of these types of proteins to be upregulated.

### Open mass searches did not reveal modifications of peptides from AHA-treated cells

A possible reason for the lack of observable AHA tags in our data is that incorporation of AHA into plant proteins causes an unexpected mass change (e.g. through posttranslational modification of the AHA moiety). To investigate this possibility, we performed open-mass searches on the data from AHA-treated and untreated cells. We used the AGIs from the enriched datasets and the HPG-tagged dataset to filter the data and then used the filtered data to construct frequency histograms for the mass modifications of peptides to look for evidence of new delta mass shifts following AHA incorporation, independent of a tight precursor mass filter for AHA. However, there was no clear evidence from the comparison of delta mass shift histograms for novel mass shifts associated with AHA-treatment of cells (Supplemental Table S7).

### Development of a high confidence set of enriched, newly synthesised proteins

From our combined data, we generated a high confidence list of 202 nascent proteins (Table 3; Supplemental Table S8) that includes proteins that were enriched by click chemistry after treatment with AHA and/or HPG and 1) found to be tagged with HPG prior to enrichment and/or 2) are known to be independently labelled with ^15^N in the same time period. Using MapMan functional categories as a guide (Schwacke et al., 2019), these proteins were classified into eight supergroups. Around one quarter of the proteins in the list were involved with protein biosynthesis. Proteins involved in nucleotide metabolism and processing, cellular organisation, nutrient uptake, photosynthesis, redox homeostasis, secondary metabolism, solute transport and vesicle trafficking and enzymes and coenzymes also featured in this list. HPG enrichment included more proteins than AHA enrichment for every supergroup of proteins examined.

**Table 3.** Functional categories of nascent Arabidopsis proteins identified by non-canonical amino acid incorporation with high confidence. This set of proteins were labelled with ^15^N and a) enriched by AHA-based BONCAT, b) enriched by HPG-based BONCAT and/or c) tagged in cell cultures after treatment with HPG. Functional categories were based on MapMan functional category assignment. Proteins were counted when they were observed to fulfil the stated criteria in at least two out of three biological replicates. Full details provided in Supplemental Table S8.

| Functional Category | No. proteins |  |  |  |
| --- | --- | --- | --- | --- |
|  | Total | AHA-enriched | HPG-enriched | HPG-tagged |
| Amino acid metabolism | 12 | 4 | 9 | 3 |
| Carbohydrate metabolism | 12 | 5 | 12 | 3 |
| Cellular respiration | 21 | 13 | 18 | 8 |
| Cellular organisation | 12 | 2 | 12 | 1 |
| Photosynthesis | 13 | 6 | 12 | 3 |
| Ribosomal subunits | 48 | 22 | 37 | 4 |
| Translation | 9 | 3 | 8 | 5 |
| Proteosome/proteolysis | 10 | 1 | 9 | 4 |
| Chaperones | 26 | 11 | 25 | 26 |
| Protein modification/translocation | 11 | 6 | 9 | 4 |
| Other | 38 | 17 | 32 | 11 |
| Not assigned | 16 | 5 | 16 | 2 |

### Post-translational modification profiles of nascent proteins differ to total protein sets

Being able to resin-enrich nascent proteins enabled us to make some important observations about the differences between nascent protein extracts and total protein extracts. We analysed the data for 10 common PTMs (see Experimental) and found that peptides of 55 % and 51 % of the proteins in total protein extracts from cell culture treated with AHA or HPG, respectively, contained one or more PTMs (Figure 6). In stark contrast, peptides from only 9 or 10 % of proteins in resin-enriched nascent protein datasets from AHA- or HPG-treated cells, respectively, contained PTMs (Figure 6). Of the 214 proteins found in both nascent and total datasets using AHA, only three (1 %) contained peptides with more PTMs in the enriched dataset than in the bulk dataset. In a similar way, of the 593 proteins found in both nascent and total datasets using HPG, only nine (2 %) contained more PTMs in the enriched dataset than the bulk dataset (Supplemental Table S9). When individual proteins were considered, the proportion of spectral hits for PTM-containing peptides per protein was greater in the bulk datasets compared to their enriched counterparts in all cases for datasets from AHA-treated cells (25 protein IDs; *p* < 0.05 (*n* = 3); Supplemental Table S9) and in all but three cases for datasets from HPG-treated cells (64 protein IDs, total; *p* < 0.05 (*n* = 3); Supplemental Table S9). Considering the larger HPG-treated dataset, the proteins that were significantly more modified in the bulk dataset compared to the nascent dataset were generally proteins related to primary metabolism or protein biosynthesis, homeostasis and modification. The high degree of modification of proteins in these pathways as they age may indicate that they are exposed for longer or to harsher (bio)chemical environments than other proteins *in vivo*.

**Figure 6.**
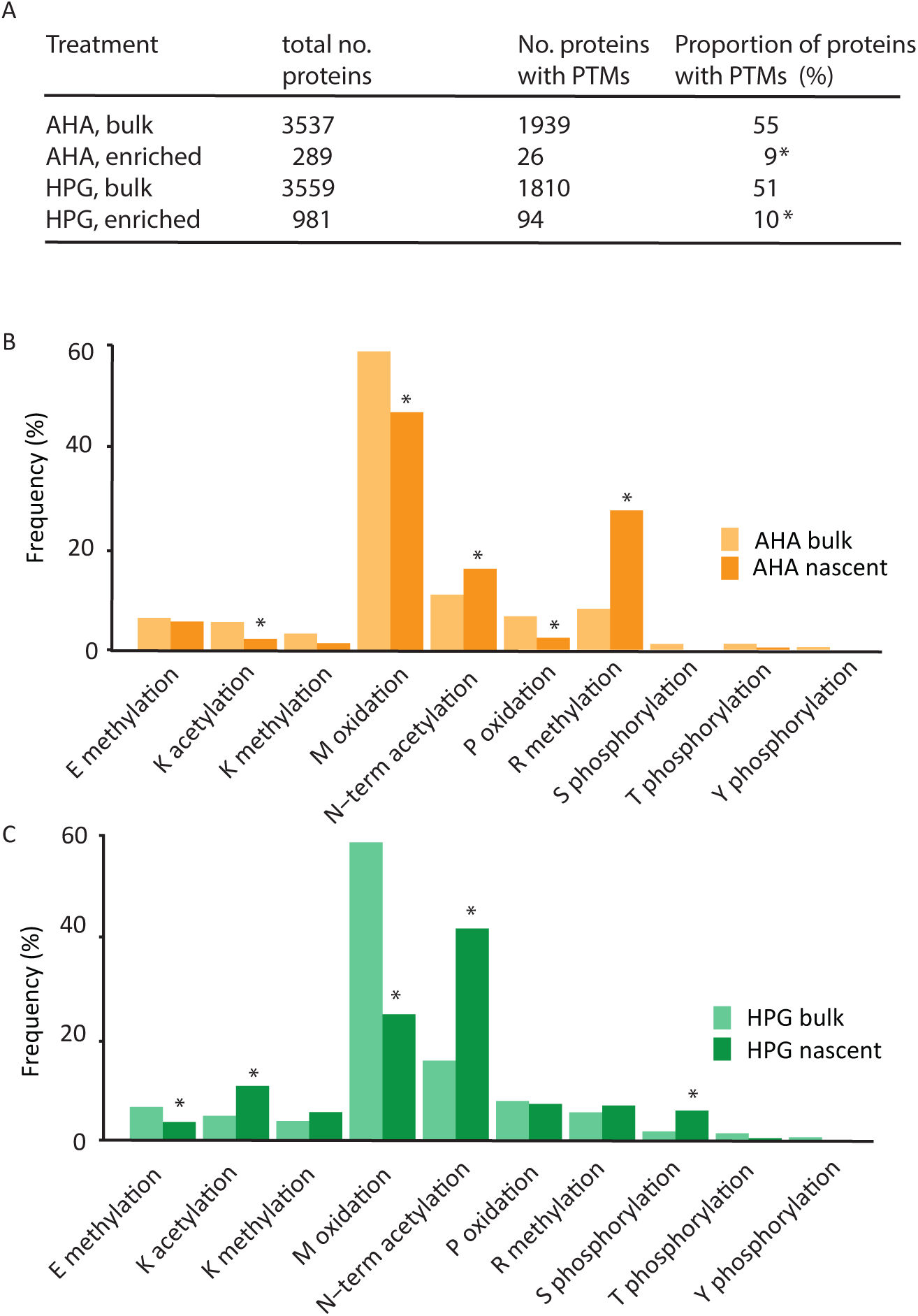
Post-translational modifications of peptides from non-canonical amino acid labelled resin-enriched and total proteins sets. Number of PTMs in ncAA enriched and total proteins sets (A), and the relative frequency of each type of modification following AHA (B) and HPG (C) tagging and enrichment. Modification frequencies were calculated using the number of spectral counts of modified and unmodified peptides. Z-tests for 2 population proportions were used to calculate p values, * denotes p<0.01.

We further analysed the data to see any differences in the relative frequency of occurrence of the ten modifications across the nascent and bulk proteins from AHA- and HPG-treated cells (Figure 6, Supplemental Table S9). This showed that the higher frequency of PTMs in the bulk protein extracts was largely due to higher frequency of Glu methylation, Met oxidation and Thr/Tyr phosphorylation.

Considering the AHA datasets, aside from small differences in oxidation (higher frequency in bulk dataset) and *N*-terminal acetylation (higher frequency in nascent dataset), the nascent and bulk proteins appeared similar in terms of PTM frequencies (Figure 6B). On first analysis it appears that there is a very high proportional frequency of Arg methylation in nascent proteins from AHA treated cell cultures (Figure 6), but, of the 129 Arg-methylated peptide spectra found in this dataset, all but two were mapped to an HSP 70 chaperone (AT3G09440.1), indicating that this is not a general difference between bulk and nascent protein sets but a case of enrichment of a specific protein with a dominant PTM.

In contrast, nascent proteins from HPG-treated cell cultures contained a much lower frequency of methionine oxidation and a much higher frequency of *N*-terminal acetylation compared to the matched bulk protein sets (Figure 6C). The *N*-terminal acetylation was found across 48 HPG-enriched proteins. The set of nascent proteins with lower number of spectral counts to PTM containing peptides included major proteins involved in the ribosome, respiratory metabolism and molecular chaperones (Supplemental Table S9).

## Discussion

### The benefit of HPG as a BONCAT agent for plants

Given the importance to find an experimental means of extracting nascent plant proteins for a wide variety of purposes, we set out to determine the utility of different BONCAT agents. We began our study in whole Arabidopsis plants, but our data showed that the previously established BONCAT agent of choice, AHA, was not efficiently incorporated into proteins in whole Arabidopsis seedlings and inhibited plant growth. We found better evidence of tagging with an alternative agent, HPG, using the same seedling flood method. Turning to Arabidopsis cell culture, we found HPG to be an efficient tagging agent, but no evidence of AHA tagging under the same conditions. Moreover, HPG tags were frequently found in the most abundant newly-synthesised proteins, as determined by independent measures of ^15^N protein labelling in matched experiments.

This difference between AHA and HPG may be due to the different efficiencies of the plant methionyl-tRNA synthetase in using AHA and HPG to charge tRNAs with these ncAAs. However, our data suggests a contributing factor for this difference in tagging effectiveness is that AHA stimulated production of Met in plant cells whereas HPG did not (Figure 2). This resulted in the level of HPG reaching 4.8 times that of Met in HPG-treated cells, but in AHA-treated cells, the level of AHA was only 1.6 times that of Met. This ratio means that AHA is less likely than HPG to be incorporated into nascent proteins due to greater Met competition in charging tRNA. Furthermore, our data show that accumulation of Met (induced by treatment with Met or AHA) in Arabidopsis cell cultures causes an elevation of 5’-methylthioadenosine, altering the balance of metabolites in the Met cycle (Figure 2). We suggest that this is a potential cause of the retarded plant and cell culture growth observed in our experiments. Our result is consistent with previous data that showed exogenous Met supplied to plant cell cultures can negatively affect cell growth (Vakkari, 1980) and the observation that the Met-over-accumulating mutant *mto2-1* has a dwarf phenotype (Kusano et al., 2010). Hence, even if AHA could efficiently tag nascent proteins, the results obtained may be confounded by the fact that AHA-treated cells are likely to be in a metabolically perturbed state and protein synthesis rate is likely to be slower overall. This perturbation of the Met cycle was not observed in cell cultures treated with HPG, which contained Met and Met-derived metabolites at levels similar to untreated plants. This feature alone makes HPG a more attractive nascent enzyme tagging agent for plants.

Even though our data showed that HPG was a preferable ncAA for BONCAT in plants compared to AHA, there is still some discrepancy between the groups of proteins that were labelled with ^15^N and those that were tagged with HPG. One reason for this is that, since Met is a relatively rare amino acid in proteins (Gilis et al., 2001), Met-to-HPG substitutions would take place far less frequently than incorporation of ^15^N, which is variably incorporated into every amino acid and thus every nascent protein. While we saw some weak correlation between protein Met content and HPG labelling (Supplemental Figure S13), in our view this was insufficient to fully explain the difference between the HPG and ^15^N datasets.

### Reconciling BONCAT data with previous reports in plants

While in our hands AHA appears to be relatively unsuitable for BONCAT due to the impact on plant and plant cell growth and the absence of AHA moieties in peptide MS data, this contrasts with other recent reports promoting the use of AHA in plants (Glenn et al., 2017; Yu et al., 2020). The approaches used, evidence provided and conclusions drawn in the current study differ from those presented by Yu et al. (2020) and Glenn et al. (2017). Yu et al. (2020) relied on the Glenn et al. (2017) report that all proteins that were enriched by the alkyne-resin linked click-chemistry were newly-synthesised. Glenn et al. (2017) used TAMRA-alkyne labelling to selectively react with azide moieties for gel based visualisation, and then LC-MS for identification of click-chemistry enriched proteins. However, our independent analysis of the PRIDE-deposited LC-MS data from Glenn et al. (2017) showed that of a total of 13229 peptides mapped to 3060 proteins, only spectra for one peptide, mapped to a plasma membrane-localized member of the RLCK-VIII subfamily (AT2G41970.1) showed the expected Met-to-AHA mass modification (−4.9863). Furthermore, this peptide was detected in the heat shock samples only and not in the control or recovery samples. Yu et al. (2020) did not deposit their raw MS data into a public repository and, upon request, were unable to provide raw data for comparison to Glenn et al. (2017) or our own data. Direct comparison of the proteins identified by Glenn et al. (2017) and Yu et al. (2020) and those identified in the enrichment sets in the present study are presented in Supplemental Figure S14. These comparisons revealed several interesting findings. Firstly, for each set of proteins identified by the four different experiments, there was substantial overlap, but also each experiment resulted in a distinct set of proteins that were not found in any other experiment (Supplemental Figure S14). This is confounded by the fact Yu et al. (2020) only provided analysis of cold-treated Col-0 and *stch4-1* mutant plants (only the results from Col-0 were used for our re-analysis). Nevertheless, even comparing our results to the controls from Glenn et al. (2017) showed only partial overlap; of the proteins enriched by HPG- and AHA-based BONCAT in the present study, 67 % and 53 %, respectively, were also identified by Glenn et al. (2017) in their control samples (Supplemental Figure S14). The most likely cause of the variation of absolute numbers of proteins identified in our study compared to those presented by Glenn et al. (2017) and Yu et al. (2020) is differences in plant materials and LC-MS analysis, rather than differences in enrichment processes per se.

The near absence of the expected Met-to-AHA mass modification in the data presented in Glenn et al. (2017) does not necessarily mean that AHA was not incorporated into nascent proteins; an alternative explanation is that, because Glenn et al. (2017) used trypsin to digest the enriched proteins while they were still bound to the enrichment resin, the AHA-tagged peptides were not present in the peptide mixture that was analysed by LC-MS. Only the peptides that were cleaved from the resin by trypsin were analysed and it is less likely that these would contain tags. There was no analysis of bulk protein samples to search for AHA tags in either previous report. Nevertheless, these results are consistent with our data from whole plants treated with AHA, which showed only one tagged peptide (not mapped to the same protein as in Glenn et al. (2017)) even in the absence of enrichment. As it was not possible to identify the TAMRA-alkyne-linked proteins in the mass spectrometry data from Glenn et al. (2017), there was no independent confirmation for the vast majority of proteins in the resin-enriched pool that they were enriched because they had been tagged with AHA. Despite these inconsistencies, a significant point of overlap between our data and the data presented by Glenn et al. (2017) is the abundant presence of heat shock proteins in both nascent protein sets.

By combining ncAA residue analysis in peptide MS data from bulk and enriched samples with quantitative protein enrichment assessments and comparison to matched ^15^N labelling studies, we suggest that multiple independent assessments should be used to confirm the utility of ncAA labelling strategies in a given plant species and sample type. While complete overlap between these datasets has not been possible, for many of the reasons outlined above, they do provide confidence that HPG tagging is a viable strategy and that enrichment, preferably confirmed by HPG residue confirmation and/or adequate evidence of ^15^N incorporation, can be used as a measure of protein synthesis and also as an extracted source of nascent proteins for further analysis. Arabidopsis protein sets that have been so confirmed in this study to be newly-synthesised by enrichment and ^15^N labelling and/or HPG tagging are summarised in Table 3 and shown in detail in Supplemental Table S8.

### Utility cases for the use of BONCAT methods in plants

Despite the technical difficulties outlined above, BONCAT with HPG appears to be a valuable new tool, for studying proteins related to protein synthesis, primary metabolism and chaperones in plants.

BONCAT could now be used to understand how components of the plant proteome respond to stimuli, not just through protein abundance changes, but by protein synthesis changes. This is especially valuable because in short timeframes, it is often difficult to detect significant changes in protein abundance. The known weak correlation between changes in mRNA levels and changes in protein abundances (Washburn et al., 2003) is likely to be underpinned by a better correlation between changes in nascent protein abundance and mRNA levels. Moreover, a range of forms of translational control are invisible to transcriptomic analysis. For example, Clark et al. (2019) showed that 1009 proteins changed in abundance in response to auxin in Arabidopsis seedlings, with no significant changes in transcript levels for any of those proteins. Therefore, assessing the metabolic consequences of plant stimuli directly through measuring rates of protein synthesis enables new types of short timeframe experiments to be undertaken. BONCAT could be used in isolation or in combination with transcriptomic analysis to give a richer understanding of plant metabolic processes in short periods of time.

BONCAT could also be used in the future to study the mechanisms of degradation of proteins. For example, it is reported that PTMs have a role in regulating protein turnover (e.g. Sridevi et al., 2009), but this is only known experimentally for a limited number of proteins. Proteins with specific protein domains tend to degrade at similar rates (Li et al., 2017), but even proteins of these sub-populations vary substantially in degradation rate. There is also a link between protein turnover rates and the age of the organism in question; as organisms grow older, protein turnover rates generally slow down (Dhondt et al., 2017; Basisty et al., 2018). Loss of ‘proteostasis’ results in proteins changing in several ways including aggregation (Kennedy et al., 2014), unfolding (Hetz et al., 2015), oxidative damage (Cannizzo et al., 2012) and increased accumulation of PTMs (Tan et al., 2007; Baskin and Taegtmeyer, 2011; Tsakiri et al., 2013; Brehm and Krüger, 2015). However, whether these changes are all results of decreased protein turnover is unclear and our understanding of the driving force for the decrease in protein turnover with age remains incomplete. In plants, mammals and yeast, the ‘*N*-end rule’ has also been reported to have a central role in controlling protein degradation rates (Bachmair and Varshavsky, 1989; Potuschak et al., 1998; Worley et al., 1998; Dissmeyer et al., 2018) but protein turnover measurements of the 1000 most abundant proteins in Arabidopsis leaves failed to reveal any correlation between turnover rates and *N*-terminal amino acids (Li et al., 2017). When we analysed our data for the presence of variable modifications we found nascent proteins sets were largely devoid of many commonly found PTMs (Supplemental Table S9), suggesting that as proteins age they accumulate PTMs. The observation that HPG-enriched proteins retain *N*-terminal acetylation in their PTM profile in comparison to other types of PTMs (Figure 6) is consistent with the co-translational nature of N-terminal protein modifications in plants (reviewed in Millar et al., 2019). The biological relevance of a lack of Met oxidation in nascent molecular chaperones and nascent mitochondrial proteins (Supplemental Table S9) can be seen in a range of reports from other organisms. For example, methionine oxidation of HSP60 has been linked to the role of this protein in defence against reactive oxygen species in human cell lines (Li et al., 2014), reversible Met oxidation of HSP70 in yeast inhibits its activity and maintains protein-folding homeostasis in suboptimal cellular folding environments (Nicklow and Sevier, 2020), and oxidised ATP synthase subunits identified here have also been reported as hot-spots of protein oxidation from mitochondrial ROS generated from the electron transport chain (Kane and Van Eyk, 2009).

Therefore, PTM accumulation may include a cumulative chemical process, through which proteins accumulate PTMs at a particular rate over time, which may well be the case for oxidation of Met and Pro residues and can have functional implications as indicated above. However, it could also be that as proteins age, they are enzymatically modified providing a marker for their age or the status of cells, and PTM pattern analysis may then enable the monitoring of the health of a protein population. Different age-dependent PTM rates may, therefore, help to explain differences in protein turnover rates that cannot be predicted from primary sequence or secondary structural features alone (Li et al., 2017) as well as impact on the functional status of the proteins themselves.

## Conclusion

BONCAT is a valuable tool for studying protein synthesis in plants, but it suffers from technical challenges that have not been encountered in other organisms, potentially associated with Met synthesis by plants. Using HPG rather than AHA appears to overcome some of these challenges. As questions remain about the selectivity of the click-chemistry method for enriching plant proteins, independent evidence from ncAA residue analysis or isotope labelling is currently needed to confirm claims. The use of BONCAT will enable new features of nascent plant proteins to be discovered, such as the accumulation of PTMs, to investigate proteins as they age *in vivo*.

## Materials and Methods

### Arabidopsis Cell Culture media

Liquid Murashige & Skoog Minimal Organics (MSMO) media was prepared as follows: sucrose (30 g; Univar), 4-morpholineethanesulfonic acid (MES; 0.5 g; Astral Scientific) and Linsmeier & Skoog Basal Medium (4.4 g; PhytoTech Labs) were dissolved in ca. 900 mL of H_2_O, α-naphthaleneacetic acid (500 µL; 1 mg/mL) and kinetin (50 µL; 1 mg/mL; Sigma) were added and the pH of the mixture was adjusted to 5.7 with aqueous KOH before dilution to 1000 mL with H_2_O. The medium was sterilised by autoclaving.

MSMO media containing ^15^N was prepared as follows: Murashige & Skoog Modified Basal Salt Mixture (0.6 g; PhytoTech Labs), ^15^NH_4_^15^NO_3_ (1.7 g), K^15^NO_3_ (1.9 g), KH_2_PO_4_ (0.2 g), *myo*-inositol (0.1 g), sucrose (30 g; Univar), and MES (0.5 g; Astral Scientific) were dissolved in ca. 900 mL of H_2_O; hormones were added, the pH was adjusted and the mixture was diluted, as described above. The medium was sterilised by autoclaving.

### Plant growth conditions

Sterilised (70 %v/v ethanol in H_2_O with 0.05 %v/v Triton X100 rotating end-over-end for 5 min before being rinsed with 100 % ethanol) *Arabidopsis thaliana* (Col-0) seeds were sprinkled onto agar plates (0.9 %w/v agar, ½ strength MS salts, 0.3 % sucrose). The seeds were vernalised fore ca. two days before being transferred into continuous light (ca. 600 µmol m^-2^ s^-1^) at 22 °C. Three-day-old seedlings were treated with AHA or HPG (1 mM in 9.9 mM potassium citrate buffer (pH 5.6, with 150 mM sucrose, 4.32 mg/mL MS salts and 0.0024 %v/v Silwett L-77) *via* seedling flood for two minutes (Glenn et al., 2017). The solutions were decanted and the plants harvested three hours later into liquid N_2_.

To determine the longer-term effects of ncAA treatment on Arabidopsis growth, the treatment procedure outlined above was repeated, except the plants were allowed to continue growing for 72 hours after treatment (in the same conditions). Two plates were subject to each treatment. Plants were photographed immediately after treatment and then again after 24, 48 and 72 h. Photographs were analysed using the ‘Threshold Colour’ plug-in in ImageJ (National Institutes of Health, 1.53a), as described by Corral et al. (2017). To control for differences in the total number of germinated seeds on each plate, the total number of green pixels in each photo were normalised as follows: the initial green areas for each replicate were averaged and a factor was generated by dividing this average by the initial green area for each replicate. Each measurement was then multiplied by this factor to generate the normalised green area.

### Measuring the effect of AHA, HPG and ^15^N on plant cell growth

Three-day-old Arabidopsis cell cultures (PBS-D) in liquid MSMO media (120 mL each) were treated with 1 mM AHA or HPG. Initial treatments were performed by adding the solid powders directly to the cell cultures to give the desired concentration. Follow-up experiments were conducted by adding enough filter-sterilised AHA or HPG solution (100 mM in H_2_O) to give final concentrations of 1 mM. Samples (each ca. ^1^/_3_ of the total culture volume) of cell cultures were harvested by vacuum filtration (Whatman No. 1 filter paper) after approximately 0, 8 and 24 hours of exposure to AHA, HPG or ^15^N. The fresh weight of each sample was recorded prior to snap-freezing in liquid N_2_. The fresh weights of a control were also obtained at these time points. Three replicates, each consisting of a separate cell culture flask, were used in each treatment. Small (< 1 mL) samples of one replicate for each treatment were also taken for viability staining using propidium iodide and fluorescein diacetate followed by microscopic analysis.

### Metabolite extraction

Cell culture samples were ground in liquid N_2_ with a mortar and pestle. Extraction solvent (80 %v/v MeOH in H_2_O, supplemented with [^13^C_5_, ^15^N]Met at 20.5 ng/µL) was added to each sample at a rate of 1 µL/mg (FW). The samples were rotated end-over-end for 17 h and then centrifuged (10 min, 2.1 x 10^4^ RCF, ambient temperature) to pellet the spent cell culture. Extracts were stored at -80 °C, until they could be analysed.

In preparation for analysis, the supernatants were transferred to clean tubes and centrifuged a second time (45 min, 1.8 x 10^4^ RCF, temperature set to 4 °C but the centrifuge was not pre-chilled). These supernatants were sub-sampled (20–200 µL) and the solvent removed under reduced pressure. The residues were re-suspended in an equal volume of 0.1 % v/v formic acid in H_2_O (except where the sub-sample was too small to allow this, in which case up to two-times the sub-sample volume was used). To ensure no particulates were present in the subsample, the re-suspensions were centrifuged (45 min, 1.8 x 10^4^ RCF, temperature set to 10 °C but the centrifuge was not pre-chilled).

### LC-MS for metabolites

LC-MS was conducted using an Agilent 6430 Triple Quadrupole LC/MS System. The LC solvents were 0.1 %v/v formic acid in H_2_O (A) and 0.1 %v/v formic acid in acetonitrile (B) (both solvents were Honeywell LC-MS Grade LabReady Blends). The flow rate was 200 µL and the gradient was as follows: 0.0 % B from 0.00 to 3.00 min, increasing steadily to 100 % B at 6.00 min; the solvent composition remained at 100 % B until 8.00 min, after which it returned to 0.0 % B over 0.50 min and the system was allowed to equilibrate at this solvent composition for 20 min. The MS was operated in positive ion mode with an ion source temperature of 125 °C, capillary voltage of 4.0 kV, a desolvation (N_2_) gas flow of 11 L/min and a nebulizer pressure of 15 psi. MRM was used to quantify all analytes (Supplemental Table S1). The cell accelerator voltage for each MRM transition was 4 V. Data was analysed using the Agilent MassHunter QQQ Quantitative Analysis software.

### Protein extraction and LC-MS preparation

Protein extraction was performed essentially according to the procedure outlined by Wessel and Flügge (1984). Briefly, extraction buffer (1.6 uL per mg (FW) of plant tissue; 125 mM Tris-HCl, pH 7.0, 7 %w/v SDS, 0.5 % PVP-40, 25 mM DTT with one cOmplete protease inhibitor tablet added per 50 mL of buffer) was added to homogenised plant tissue (ca. 250 mg (FW)) and, after vortexing, the solid was removed by centrifugation; this also removed some of the liquid, leaving 200 uL per sample, to which was added methanol (800 uL), chloroform (200 µL) and H_2_O (500 µL) and the resulting emulsion was split by centrifugation. The protein pellet was washed twice with both ice-cold methanol and acetone (one acetone wash was left for one hour at -20 °C and the subsequent one was left for 12– 17 h at -20 °C. The protein pellets were re-suspended in NH_4_HCO_3_ (50 mM) with SDS (1 %w/v) and DTT (10 mM) in H_2_O. Protein concentration was determined using Amido Black assays. Disulfide bonds were reduced with iodoacetamide and the proteins were digested using trypsin. Digested protein samples were fractionated using HPLC (a J4-SDS column mounted to a C_18_ analytical column) into 96 (12 columns x 8 rows) and these fractions were combined by column into groups of eight to give 12 fractions. Clean peptide fractions were dried under reduced pressure and stored at -80 or -20 °C until they could be analysed.

### Click chemistry enrichment of AHA/HPG tagged proteins

For experiments where enrichment of azide-(AHA) or alkyne-(HPG) tagged proteins was required, Click Chemistry Tools Click-&-Go Protein Enrichment Kits (Product numbers 1152 and 1153) were used, with minimal modifications to the protocol. Tissue samples (1– 2 g) were extracted as described above (except that the precipitation and washing step volumes were increased seven-fold) up to completion of the acetone wash steps. After this point, the pellets were re-suspended in 3.2 mL of Lysis Buffer and the protein concentrations determined using Amido Black assays. The click reaction was performed as described in the protocol, except the reaction mixtures were prepared as follows: protein solution: 2400 uL (containing 1 mg of protein), 2X Cu catalyst solution: 3000 uL, resin: 200 uL, H_2_O: 400 uL. After the resin-bound proteins were washed, the resins were each split approximately in half and one half of each was treated with trypsin and the other was treated with hydrazine hydrate (2 %v/v solution) and then washed with phosphate-buffered saline (containing 1 %w/v SDS, as recommended by the supplier, although the protocol does not call for SDS in this solution) to cleave the bound proteins from the resin. The resin that had been treated with trypsin was then also treated with hydrazine hydrate (2 %v/v solution) to cleave the peptides that remained bound to the resin after digestion. The proteins that had been cleaved from the resin with hydrazine were precipitated using ice-cold acetone (≥ 10x solution volume). The resulting protein pellets were resuspended in digestion buffer (according to the protocol) and digested with trypsin. The enriched samples that were cleaved from the resin (both those that were cleaved off as whole proteins and those that were cleaved after trypsin digestion) were not fractionated, rather residual SDS and salt was removed by HPLC using a J4-SDS column mounted to a C_18_ trap. These peptide samples were purified further, along with the peptides that had been generated by digestion of the resin-bound proteins, using silica C_18_ MacroSpin Columns (Nest Group). Clean peptide fractions were dried under reduced pressure and stored at -80 or -20 °C until they could be analysed.

### Peptide mass spectrometry

Clean and dry peptide samples were re-suspended in MeCN/H_2_O (2 %v/v) with formic acid (0. 1 %v/v) and filtered through 0.22 µm filters (Millipore) before being loaded for LC separation. A total of 4 µL of each filtered sample was loaded onto a C_18_ high-capacity nanoLCchip (Agilent Technologies) using a 1200 series capillary pump (Agilent Technologies) as described previously. Following loading, samples were eluted from the C_18_ column directly into a 6550 series QToF MS (Agilent Technologies) with a 1200 series nano pump using the following gradient. B (0.1% formic acid in acetonitrile) gradient: 100 to 3 % B in 1 min, 3 to 35 % B in 55 min, 35 to 95% in 3 min, and 95 to 3 % in 2 min; the system was allowed to re-equilibrate for 15 min at this solvent composition before the next run. Parameter settings in the mass spectrometer and data acquisition settings were as described previously (Nelson et al., 2014), except the ions were excluded for 0.12 min (rather than 0.4 min) following fragmentation and the minimum threshold for precursor ion selection was 5,000 counts (instead of 10,000 counts). The MS proteomics data have been deposited to the ProteomeXchange Consortium via the PRIDE partner repository (Perez-Riverol et al., 2019) with the dataset identifier PXD023170.

### Analysis of peptide mass spectra

For ^15^N label incorporation analysis, Agilent .d files were converted to mzML using the Msconvert package (v3.0.1908-43e675997). These files were analysed using the Trans Proteomic Pipeline (a joint project between the Institute for Systems Biology and the Seattle Proteome Center) and then using an in-house R-script (see Li et al., 2017).

To detect AHA and HPG tagging, Agilent .d files were analysed using MaxQuant (MQ, v1.6.10.43; Cox and Mann, 2008), with Met-to-AHA (−4.9863 Da), Met-to-HPG (−21.9877 Da) and Met-to-triazole (+50.0559 Da) as additional variable modifications. MQ output files from tagging experiments and the untreated control were processed using in-house R-scripts (available from https://github.com/ndtivendale/ncaa).

PCAs was performed using the ‘ggbiplot’ package. Only proteins that riBAQ values greater than 10^-5^ were used for PCA.

GO term enrichment analysis was conducted using AgriGO (Du et al., 2010; Tian et al., 2017), with the default parameters. The background list of protein groups included all groups detected in all treatments and experiments reported in this study. Four foreground lists were used: ^15^N-labelled, HPG-tagged, present after treatment with AHA followed by enrichment and present after treatment with HPG followed by enrichment.

GO and PO domain enrichment analysis was conducted in R using in-house scripts (available from https://github.com/ndtivendale/ncaa). The background for the HPG tagging and enrichment was the total set of proteins from bulk samples for HPG-treated cells. Similarly, the background for the AHA enrichment was the total set of proteins from bulk samples for AHA-treated cells.

For open searches, one complete replicate of AHA, HPG and untagged raw data was searched against the Araport11 peptide database with a precursor mass tolerance of ± 250 amu using Comet (2019.01 rev5). Results were rescored using Peptide prophet (TPP5.0), filtered to 1% FDR and histograms of mass error prepared using R.

For detection of modifications on enriched and bulk proteomics datasets from AHA and HPG tagging experiments, MSFragger run in the FragPipe wrapper was used (Kong et al., 2017). In addition to the Met-to-AHA and Met-to-HPG variable modifications and the carbamidomethylation of Cys fixed modification, the following variable modifications were allowed: Met or Pro oxidation (+15.9949 Da), *N*-terminal or Lys acetylation (+42.0106 Da), Ser, Thr or Tyr phosphorylation (+79.9663 Da) and Glu, Arg or Lys methylation (+14.0157 Da). Only one modification per peptide was allowed. Results were checked using the Philosopher program (da Veiga Leprevost et al., 2020) and data was analysed using an in-house R-script (available from https://github.com/ndtivendale/ncaa). To pass the filtration steps of this script, a protein had to be identified in two out of three replicates and a protein was considered modified if it was identified as modified in two out of three replicates. If a protein was found to be modified in only one out of three replicates, it was deemed to have an unknown modification status and removed from further analysis steps. When comparing the extent of modification between the enriched and bulk samples, Met-to-AHA, Met-to-HPG and Cys carbamidomethylation were excluded. For t-tests to detect significant differences between the enriched and bulk datasets in the modified:total spectral hits for each protein, we only considered proteins identified with a minimum of five spectral hits in both the enriched and bulk samples. Z-tests of two population proportions were used to detect significant differences between proportions of proteins with PTMs and the proportion of different PTMs between bulk and resin-enriched sets.

## Supplemental Data

**Supplemental Table S1.** Details of MRM transitions for quantifying seven metabolites from Arabidopsis cell culture extracts.

**Supplemental Table S2.** Relative quantification of specific protein groups from AHA-treated Arabidopsis cell cultures.

**Supplemental Table S3.** Relative quantification of specific protein groups from HPG-treated Arabidopsis cell cultures.

**Supplemental Table S4.** Relative enrichment of specific protein groups from AHA-treated Arabidopsis cell cultures by alkyne functionalised resin.

**Supplemental Table S5.** Relative enrichment of specific protein groups from HPG-treated Arabidopsis cell cultures by azide functionalised resin.

**Supplemental Table S6.** Evidence for new protein synthesis of specific protein groups from Arabidopsis cell culture after ^15^N labelling, HPG or AHA tagging and or azide/alkyne enrichment.

**Supplemental Table S7.** Open mass searches of Met containing peptides of resin-enriched proteins from AHA treated cells that were previously shown to be experimentally tagged by HPG.

**Supplemental Table S8.** High confidence set of 202 newly synthesised proteins identified by 15N labelling and AHA or HPG treated, tagging and/or enrichment with azide or alkyne resin

**Supplemental Table S9.** The number and type of posttranslational modified vs unmodified spectra counts linked to specific protein groups in HPG and AHA-treated cell culture and in azide and alkyne resin enriched protein samples from the same.

**Supplemental Table S10.** Comparison of protein groups from alkyne and azide enriched datasets from AHA and HPG treatments in this work with those reported by Glenn et al. (2017) and Yu et al. (2020) from AHA treatment and alkyne resin enrichment.

## Author Contributions and Acknowledgements

The work was supported by funding from the Australian Research Council (CE140100008; DP180104136) to AHM. NDT and AHM designed the research. NDT and RF performed the research. NDT and OD analysed the data. NDT and AHM wrote the paper.

